## Supplementary figures and tables for "Smart 3D super-resolution microscopy reveals the architecture of the RNA scaffold in a nuclear body"

**This PDF file includes:**

Supplementary notes for videos 1 - 5

Supplementary Figures 1 - 7

Supplementary Tables 1 - 8

**Supplemental notes for videos**

Supplemental Video 1: Shell-stained paraspeckles (bounding box sorting)

- This video shows all rotationally and translationally aligned paraspeckle images, sorted by their length as calculated from the 50% intensity drop-off point. On the left, we show a 3D reconstruction composed of 5 isosurfaces from 50% to 99% of the maximum image intensity. On the right, we show the 3 cross-sections through the image center.

Supplemental Video 2: Averaged Shell-stained paraspeckles (bounding box sorting)

- This video shows the bin-averaged paraspeckles. On the left, we show a 3D reconstruction composed of 5 isosurfaces from 50% to 99% of the maximum image intensity. On the right, we show the 3 cross-sections through the image center.

Supplemental Video 3: Averaged Shell-stained out of ROI paraspeckles (covariance sorting)

- This video shows the bin-averaged averaged out-of-ROI paraspeckles. On the left, we show a 3D reconstruction composed of 5 isosurfaces from 50% to 99% of the maximum image intensity. On the right, we show the 3 cross-sections through the image center.

Supplemental Video 4: showing all 3 /5' stained paraspeckles with ellipticity ≤ 1.3. Green: 5' signal; magenta: 3' signal.

- This video shows isosurfaces of the signal from single spherical paraspeckles (ellipticity ≤ 1.3) stained at the 5'-end (green) and the 3'-end (magenta) of *NEAT1_2*. The paraspeckles were rotated to align the polarization vector with the vertical axis of the image frame and the images were sorted by ascending degree of polarization. The degree of polarization is indicated above each isosurface.

Supplemental Video 5: showing all 3 /5' stained paraspeckles with ellipticity > 1.3. Green: 5' signal; magenta: 3' signal.

- This video shows isosurfaces of the signal from single elongated paraspeckles (ellipticity > 1.3) stained at the 5'-end (green) and the 3'-end (magenta) of *NEAT1_2*. The paraspeckles were rotated to align the polarization vector with the vertical axis of the image frame and the images were sorted by ascending degree of polarization. The degree of polarization is indicated above each isosurface.

### Supplementary Figures

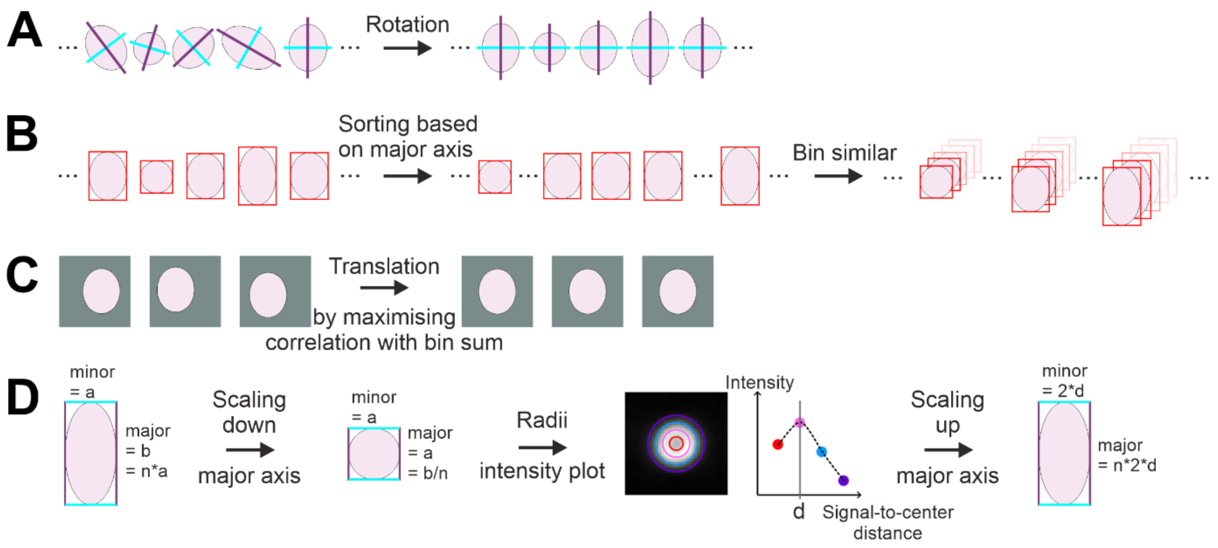


**Figure S1. Schematics of the image processing steps.** 2D schematic depictions of the different processing steps after the image acquisition. In reality, the images are 3D, but the same principles apply. **A** The raw images are rotated to align the paraspeckle's major axis with the vertical image axis (**Methods**). **B** The rotationally aligned images are sorted based on their major axis length, obtained using a bounding box (**Methods**). Images with a similar major axis length are binned together. **C** Images within each bin are translated to maximize their cross-correlation with the sum over all images in that bin. **D** The major axis of each image is then scaled down to match its minor axis and a radial intensity plot is used to determine the average signal-to-center distance of the paraspeckle along the minor axis. The major axis length is then determined by multiplying this minor axis length with the major/minor axis scaling factor. A more detailed explanation of image processing is given in the **methods**.

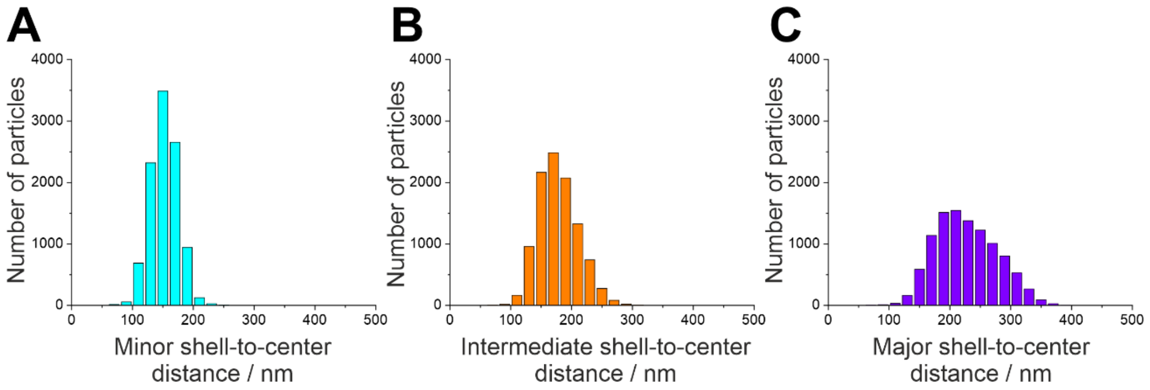


**Figure S2. Distribution of the paraspeckle lengths along the three axes.** Frequency histograms of paraspeckle lengths along their **A** minor **B** intermediate and **C** major shell-to-center distances. N=13,801 paraspeckle containing volumes from 34 independent experiments (**Table 2**).

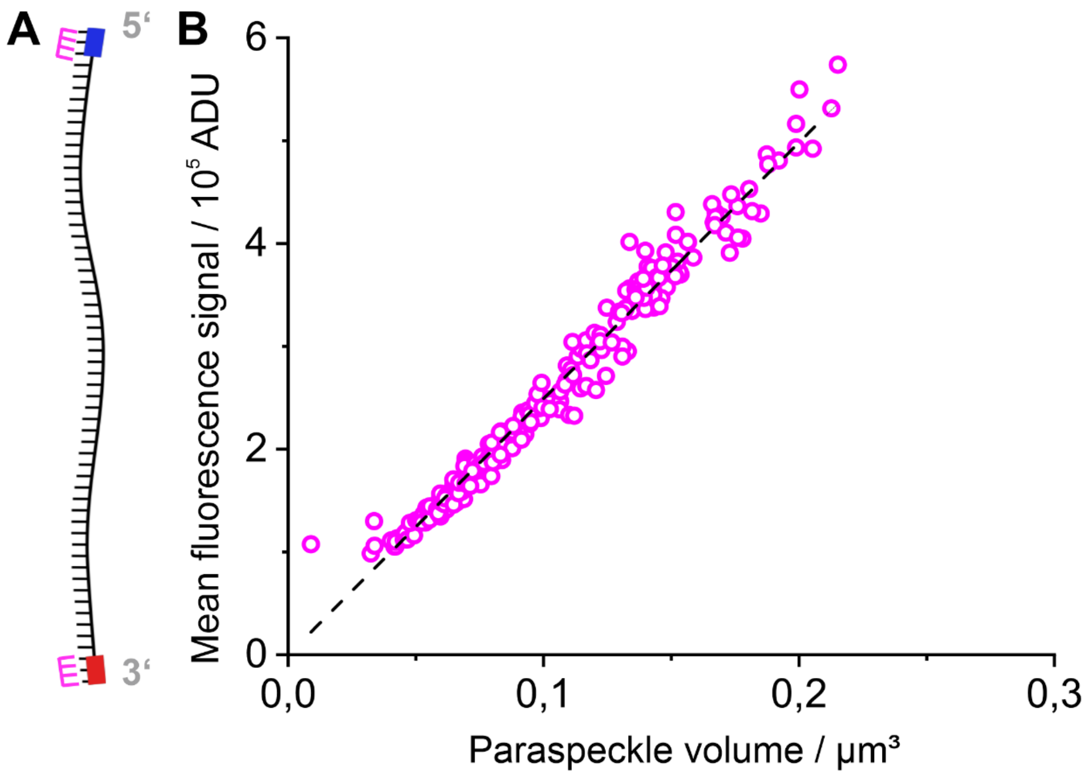


**Figure S3. Fluorescence intensity of reference probes scales with Volume.** (**A**) Scheme of RNA-FISH staining at the shell reference positions at the ends of *NEAT1_2*. (**B**) Total fluorescence intensity plotted as a function of paraspeckle volume for the shell reference. Each point represents one bin of similar paraspeckle images (see **S1B**) and is calculated by averaging the summed image intensities in that bin. N=13,801 paraspeckle containing volumes from 34 independent experiments (**Table 2**). Fit results for linear regression (black dotted line) of **B** is given in **Table 4**.

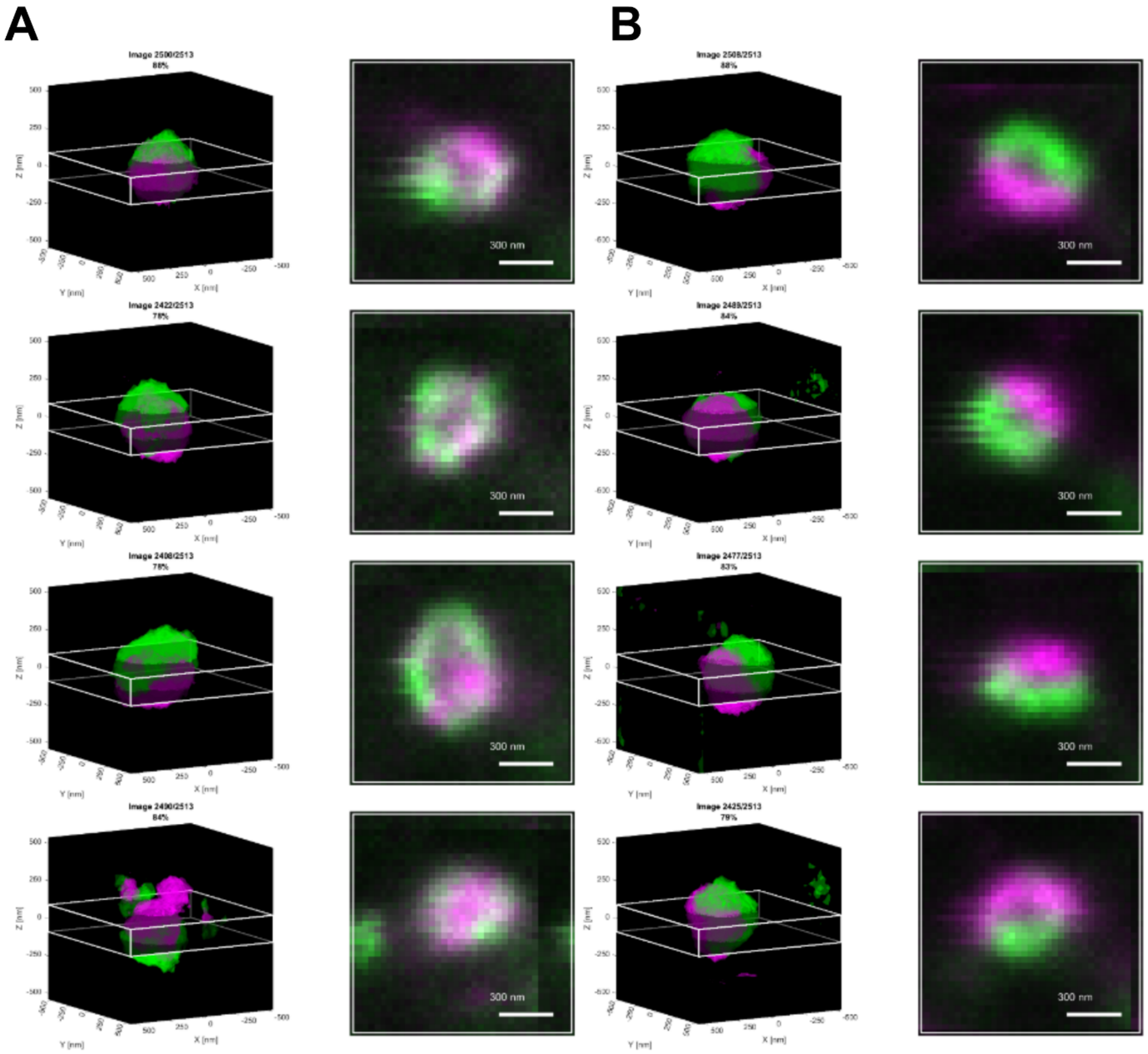


**Figure S4. Representative isosurface plots and corresponding 2D projections of paraspeckle volumes with different polarizations.** Selected paraspeckles with a degree of polarization above 78%**. A** Four examples of paraspeckles for which their polar nature is not apparent from the 2D projections. **B** Four examples of paraspeckles for which their polar nature is identifiable in 2D as well as in 3D. *2D projections correspond to 200 nm thick slices; Scale bar: 300 nm. Green: 5' signal; magenta: 3' signal.*

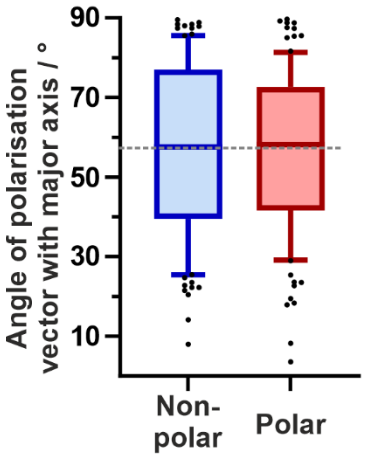


**Figure S5. Angle of polarization vector relative to the major axis for nearly spherical paraspeckles.** Angle for spherical paraspeckles (elongation ≤1.3), grouped into non-polar (100 paraspeckles with lowest polarization, 3 independent experiments, blue) and polar (100 paraspeckles with highest polarization, 3 independent experiments, red). The horizontal gray dotted line indicates the random angle of 57°.

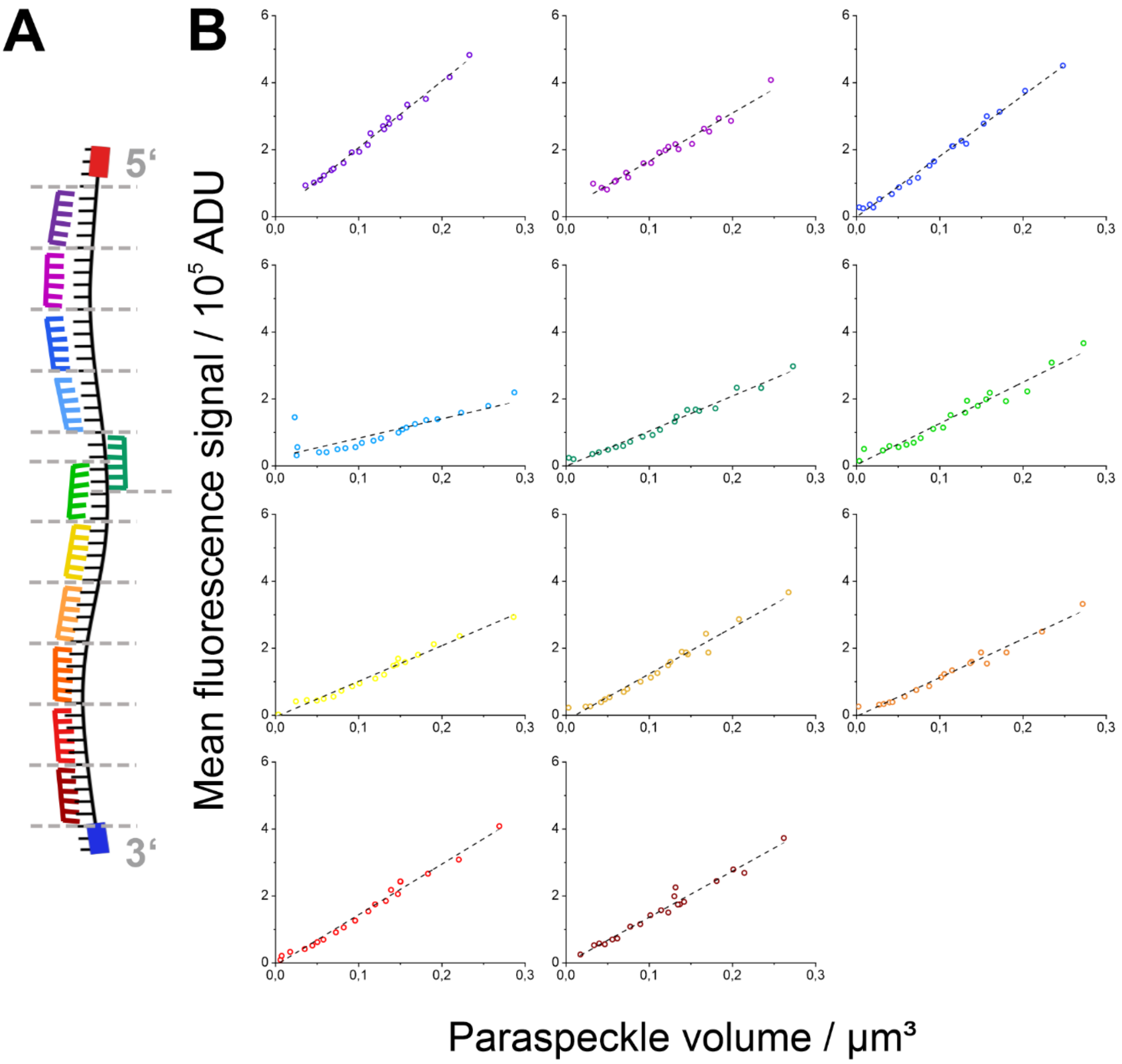


**Figure S6. Fluorescence intensity of internal probes scales with Volume.** (**A**) Scheme of RNA-FISH staining at the different 2K nt regions along the *NEAT1_2* RNA. (**B**) Total fluorescence intensity plotted as a function of paraspeckle volume for the different regions along *NEAT1_2*. Each point represents one bin of similar paraspeckle images (see **S1B**) and is calculated by averaging the summed image intensities in that bin. N=967-2992 paraspeckle containing volumes from up to 4 independent experiments were performed per *NEAT1_2* region (**Table 2**). Fit results for linear regression (black dotted lines) of **B** is given in **Table 4**.

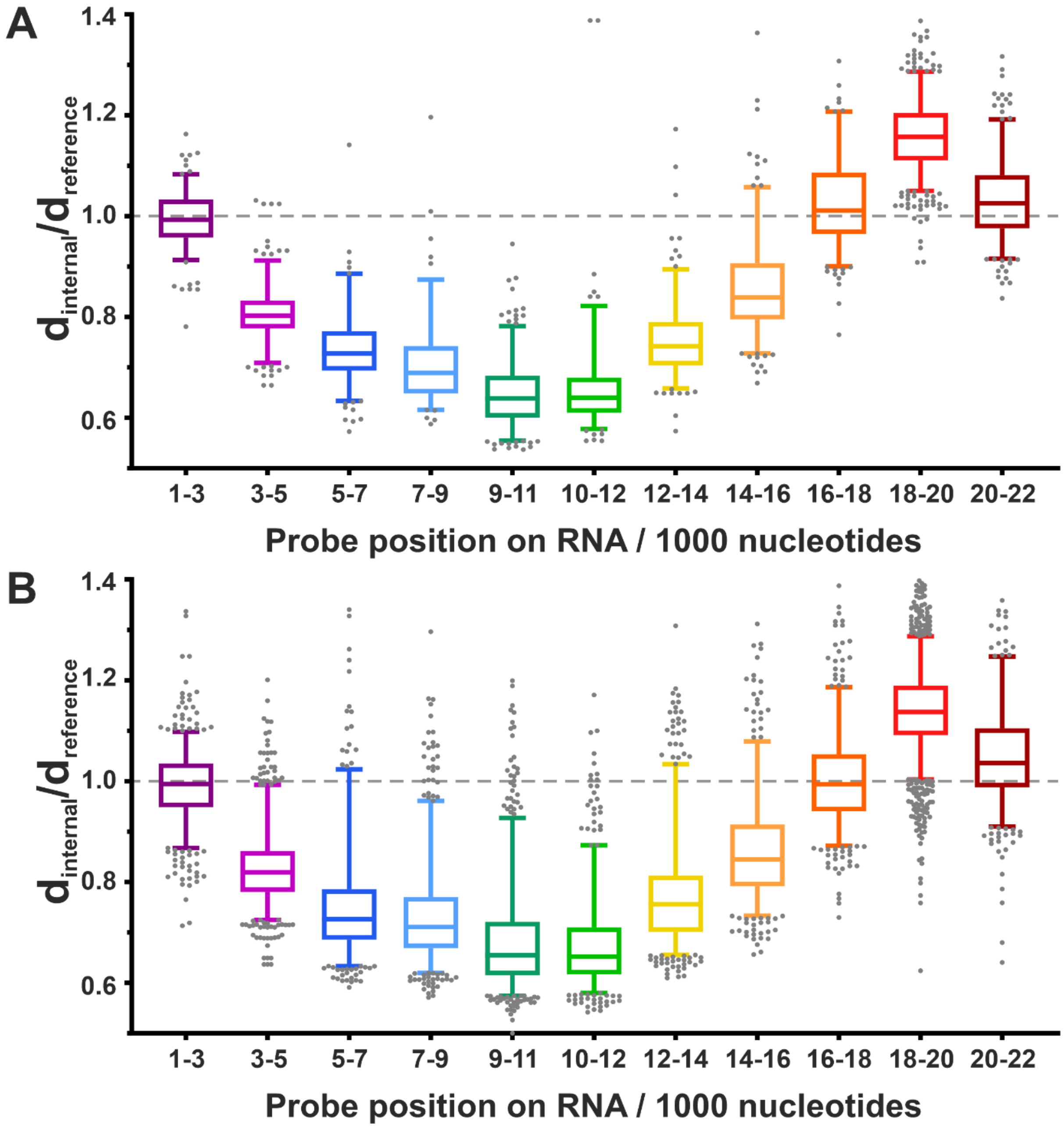


**Figure S7.** The average distance of the 11 different *NEAT1_2* sections to the paraspeckle center, relative to the to-center distance of the combined 5'/3' reference positions, as calculated from the individual images' radial intensity distribution. Results are shown for spherical paraspeckles (ellipticity ≤ 1.3) (**A**) and for ellipsoidal (ellipticity > 1.3) (**B**). N=967-2,992 paraspeckle containing volumes from up to 4 independent experiments were performed per *NEAT1_2* region (**Table 2**).

### Supplementary Tables

*Table 1: Imager and docking strand sequences used in this study. Modified LNA nucleotides are colored blue[1]. Docking sequences were synthesized together with the FISH targeting sequences from 5' to 3' end.*

| Name | Sequence | Supplier |
|---|---|---|
| P2-STAR 635P | 5' TATG**TAG**ATAA-STAR 635P-3' | biomers.net |
| P3-STAR 580 | 5' TAA**TGA**AGAAA -STAR 580-3' | biomers.net |
| P2 docking strand (FISH probe) | 5' *FISH probe sequence*-TTATCTACATA-3' | Eurofins Genomics |
| P3 docking strand (FISH probe) | 5' *FISH probe sequence*-TTTCTTCATTA-3' | Eurofins Genomics |

*Table 2: FISH staining schemes and number of recorded volumes used for figures presented in this study. For the respective figure, the corresponding section is indicated which was marked, followed by the connected docking strand. For the microscopy measurement, the corresponding imager strand was added to the imaging buffer.*

| Experiment (figure) | Staining scheme of nucleotide position (magenta = P3-STAR 580; green = P2-STAR 635P) | Number of images recorded | Number of independent measurements |
|---|---|---|---|
| Fig2B | 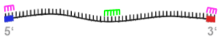 | 1 | 1 |
| Fig3, S2, S3 | 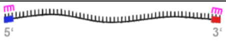<br>Reference(0-1K; 22-22.7K)-P3 | 13801 | 34 |
| Fig4, S5 | 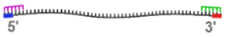<br>5prime(0-4K)-P3<br>3prime(16-22K)-P2 | 2513 | 3 |
| Fig5, S6, S7 | 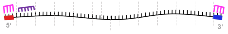<br>Reference(0-1K; 22-22.7K)-P3<br>Sliding(1-3K)-P2 | 1095 | 3 |
| | 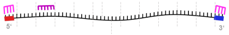<br>Reference(0-1K; 22-22.7K)-P3<br>Sliding(3-5K)-P2 | 1326 | 4 |
| | 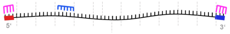<br>Reference(0-1K; 22-22.7K)-P3<br>Sliding(5-7K)-P2 | 973 | 3 |
| | 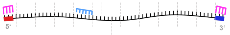<br>Reference(0-1K; 22-22.7K)-P3<br>Sliding(7-9K)-P2 | 967 | 3 |
| | 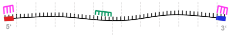<br>Reference(0-1K; 22-22.7K)-P3<br>Sliding(9-11K)-P2 | 1335 | 2 |
| | 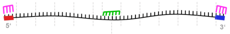<br>Reference(0-1K; 22-22.7K)-P3<br>Sliding(10-12K)-P2 | 1029 | 3 |
| | 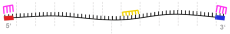<br>Reference(0-1K; 22-22.7K)-P3<br>Sliding(12-14K)-P2 | 1048 | 1 |
| | 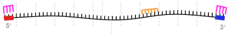<br>Reference(0-1K; 22-22.7K)-P3<br>Sliding(14-16K)-P2 | 1020 | 3 |
| | 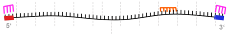<br>Reference(0-1K; 22-22.7K)-P3<br>Sliding(16-18K)-P2 | 1018 | 3 |
| | 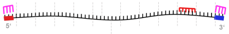<br>Reference(0-1K; 22-22.7K)-P3<br>Sliding(18-20K)-P2 | 2992 | 3 |
| | 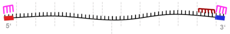<br>Reference(0-1K; 22-22.7K)-P3<br>Sliding(20-22K)-P2 | 999 | 3 |

*Table 3: Shell-to-center distances of paraspeckles measured from single-color STED measurements. The errors given are the s.e.m.*

| Ellipticity | Major axis / nm | Intermediate axis / nm | Minor axis / nm |
|---|---|---|---|
| 1.14 ± 0.9 | 126 ± 18 | 133 ± 21 | 114 ± 18 |
| 1.30 ± 0.15 | 187 ± 11 | 163 ± 20 | 146 ± 14 |
| 1.49 ± 0.14 | 234 ± 10 | 186 ± 23 | 160 ± 16 |
| 2.0 ± 0.3 | 315 ± 25 | 215 ± 32 | 162 ± 16 |

*Table 4: Values for linear regression of fluorescence intensities measured for targeted nucleotide positions of NEAT1_2 RNA. Errors are given as standard error.*

| Target nucleotide position (nucleotides form 5' to 3') / 1000 nucleotides | Slope / $10^5$*ADU/µm$^3$ | Intersection with the Y-axis / $10^5$*ADU | $R^2$ |
|---|---|---|---|
| 0-1 and 22-22.7 | 24.9 ± 0.3 | -0.01 ± 0.04 | 0.9746 |
| 1-3 | 19.9 ± 0.5 | 0.07 ± 0.06 | 0.99067 |
| 3-5 | 14.4 ± 0.6 | 0.22 ± 0.08 | 0.96993 |
| 5-7 | 18.1 ± 0.4 | -0.01 ± 0.05 | 0.99273 |
| 7-9 | 5.7 ± 0.9 | 0.30 ± 0.14 | 0.70014 |
| 9-11 | 10.5 ± 0.4 | -0.01 ± 0.05 | 0.9780 |
| 10-12 | 12.3 ± 0.7 | 0.03 ± 0.09 | 0.95826 |
| 12-14 | 10.6 ± 0.4 | -0.05 ± 0.05 | 0.98243 |
| 14-16 | 13.9 ± 0.6 | -0.15 ± 0.07 | 0.97511 |
| 16-18 | 11.6 ± 0.5 | -0.03 ± 0.06 | 0.97736 |
| 18-20 | 15.1± 0.5 | -0.07 ± 0.06 | 0.98644 |
| 20-22 | 13.8 ± 0.6 | -0.01 ± 0.08 | 0.97211 |

*Table 5: Degree of polarization (P) of paraspeckles measured from two-color STED measurements. The errors given are the s.e.m.*

| Ellipticity | Degree of Polarization (P) / % |
|---|---|
| ≤1.3 | 44.1 ± 0.8 |
| >1.3 | 39.8 ± 0.6 |

*Table 6: Angle between polarization vector and major axis (α) of paraspeckles measured from two-color STED*

*measurements. The errors given are the s.e.m.*

| Ellipticity | Degree of Polarization (P) / % | Angle between polarization vector and major axis (α) / ° |
|---|---|---|
| ≤1.3 | <14.7 | 57 ± 3 |
| ≤1.3 | >72.0 | 57 ± 3 |
| >1.3 | <8.3 | 59 ± 3 |
| >1.3 | >74.1 | 72 ± 2 |

*Table 7: Ratio of internal to reference distance to the center (d) for the 11 different internal probes measured.*

| Target nucleotide position (nucleotides form 5' to 3') / 1000 nucleotides | Ellipticity | Median d(internal)/d(reference) | 5% Percentile d(internal)/d(reference) | 95% Percentile d(internal)/d(reference) |
|---|---|---|---|---|
| 1-3 | all | 0.9943 | 0.8713 | 1.093 |
| 3-5 | all | 0.8137 | 0.7248 | 0.9811 |
| 5-7 | all | 0.7270 | 0.6333 | 1.005 |
| 7-9 | all | 0.7076 | 0.6195 | 0.9516 |
| 9-11 | all | 0.6474 | 0.5704 | 0.8863 |
| 10-12 | all | 0.6500 | 0.5800 | 0.8541 |
| 12-14 | all | 0.7502 | 0.6569 | 1.011 |
| 14-16 | all | 0.8435 | 0.7327 | 1.073 |
| 16-18 | all | 0.9980 | 0.8829 | 1.189 |
| 18-20 | all | 1.143 | 1.019 | 1.286 |
| 20-22 | all | 1.033 | 0.9142 | 1.233 |
| 1-3 | ≤1.3 | 0.9927 | 0.9129 | 1.083 |
| 3-5 | ≤1.3 | 0.8024 | 0.7090 | 0.9121 |
| 5-7 | ≤1.3 | 0.7275 | 0.6336 | 0.8860 |
| 7-9 | ≤1.3 | 0.6890 | 0.6159 | 0.8741 |
| 9-11 | ≤1.3 | 0.6387 | 0.5548 | 0.7818 |
| 10-12 | ≤1.3 | 0.6397 | 0.5779 | 0.8217 |

| | | | | |
|---|---|---|---|---|
| 12-14 | ≤1.3 | 0.7419 | 0.6585 | 0.8944 |
| 14-16 | ≤1.3 | 0.8387 | 0.7279 | 1.057 |
| 16-18 | ≤1.3 | 1.011 | 0.9004 | 1.207 |
| 18-20 | ≤1.3 | 1.157 | 1.050 | 1.286 |
| 20-22 | ≤1.3 | 1.026 | 0.9154 | 1.192 |
| 1-3 | >1.3 | 0.9943 | 0.8677 | 1.098 |
| 3-5 | >1.3 | 0.8192 | 0.7251 | 0.9928 |
| 5-7 | >1.3 | 0.7264 | 0.6333 | 1.023 |
| 7-9 | >1.3 | 0.7108 | 0.6198 | 0.9608 |
| 9-11 | >1.3 | 0.6547 | 0.5746 | 0.9273 |
| 10-12 | >1.3 | 0.6521 | 0.5801 | 0.8729 |
| 12-14 | >1.3 | 0.7560 | 0.6555 | 1.034 |
| 14-16 | >1.3 | 0.8450 | 0.7331 | 1.079 |
| 16-18 | >1.3 | 0.9939 | 0.8719 | 1.186 |
| 18-20 | >1.3 | 1.137 | 1.004 | 1.287 |
| 20-22 | >1.3 | 1.036 | 0.9101 | 1.247 |

*Table 8: List of FISH probes used to stain Neat1_2 RNA in this study.*

| Target nucleotide position (nucleotides form 5' to 3') | Seqences (5' to 3') |
|---|---|
| Reference (0-1K; 22-22.7K) | tgcggatattttccatgcag<br>ctcgctcagctatgcaagag<br>caagttgaagattagccctc<br>agcccttggtctggaaaaaa<br>aagttcagttccacaagacc<br>caggccgagcgaaaattaca<br>ttaactccacatcactcctc<br>ctgtcaaacatgctaggtgc<br>aagcgttggtcaatgttgtc<br>aaaaggagcactgccacctg<br>gtggagtgagctcacaagaa<br>cttaccagatgaccaggtaa<br>acaataccgactccaacagc<br><br>actgccatcactcgaaatga<br>tccattctctgatctgtcag<br>agtggaaatggttctctggg |

| | |
|---|---|
| | ctaagacctgtggacacttc<br>tcttttcatctgaacaggga<br>tttttgactctatgcagatc<br>cttgcattttatacccatgg<br>gaggacatctctgtgtagta<br>atttccagagtgagtctagg<br>acgagcagaggcacaagctt<br>cacaggtgagaaggtgctgg<br>aactgaacacgaggcacagt<br>ccaaggagcatgaagtcaga<br>cagcttcacatccacatgtc<br>taaaggcatagccaggggac<br>aaaagaaacacctgcggcgg<br>aaaacctgagtgcggccatg<br>ttcacagatgggaaggtgaa<br>tttgctttttcctgctcact |
| 1-3K | tcctcttactagaatgccaa<br>actgtatctctaaccaaccc<br>aattcccttcaacctgcatt<br>gcaacaacttaaccaacact<br>ctaagcaacttctcacttcc<br>taacacttcttcagtcttcc<br>actctgtgtcacctgttttc<br>cctttggttctcggaaaact<br>tgtgagatggcatcacacac<br>ccaggaggaagctggtaaag<br>ctctgaaacaggctgtcttg<br>cacttgataacacccacacc<br>cccatctttcaagtgactaa<br>cagcgaaggatgctgatctg<br>aaccacctaagttgctaagg<br>cacaccttcacagtttcaaa<br>gtggtcccttaaatacgtta<br>agaagagcccatctaatctc<br>acacctgtgacaaatgagga<br>gatgtgtttctaaggcacga<br>tgccattattaatatcgacc |
| 3-5K | tacagtgaccacaaaaggtt<br>gcaaaggtacatggattctg<br>attttaggtgatagtttccc<br>tttttaggtcttgttttcca<br>catgtagtaaaggcacctcg<br>ccattggtattactttacca<br>ggttaagaactgaactacca<br>ttgtttgcatcatccccaag<br>ctctaaatcccaacgacagt<br>cccatacatgcgtgactaat<br>atttcacaacagcatacccg<br>ccagtactttcaaccatcta<br>agttcttaccatacagagca<br>ttgtgctgtaaaggggaaga |

| | |
|---|---|
| | tgataggggtcgagaaatgt<br>ccaaagtcgttatgaaggca<br>ggagctagcaaatctagacc<br>tcctacatggccttaattac<br>ttctatctgtggttaccatg<br>gcagaccacttgggataata<br>accaactctcaacagagagc<br>cttgagtggtttcatcacat<br><br>aaagtactccccacctacac<br>accacttagaactctaacca<br>cacttagacccaaatcccag<br>agtgaggtaagaattacgca<br>aagcccagacgttttagtca<br>ttactgtagcacgcacacag<br>caagatgtatcctccagatt<br>cagtgcaaggacagaacagt<br>cccaagaggtaacacagaac<br>tcctgtaagcaaaaccttac<br>cattctacgcccaaagtttg<br>aaacaagcaccttccttctc<br>caggtgagtagaacaagcct<br>agacacagttgcaaggacgg<br>ggtcaccaagatataacggc<br>tttgtcagtagtgtacacgt<br>ccatgagcacagacagaagg<br>aaagaggaccctggattctg<br>cagggaaaggagttgcaagc<br>ataggaaaggaggcctggat<br>gagaaggcggttatagaggt<br>caggacgagaaacaaggagt |
| 5-7K | cagggtggaagcaagagttt<br>ggcatctatagcagttatct<br>tcaattcttcttcagcatgc<br>caatccccaacatttatacc<br>tctgctcctgtatcaacaag<br>attggagtggaggcaaggag<br>tgaaggcaaagaatgtggcc<br>tctcacctgatctgaggata<br>taattgctggcactgccatc<br>aaggatgcaaacggattagc<br>gtctgaattcagatctatgc<br>cacacctctggaaattcaca<br>cccactcactgaattctatt<br>acttttgctattcttatgcc<br>actgaagagcctcaaccaat<br>tcatagtcagtcagatgcac<br>ctggcattaagggtccttgg<br>actgcacatggagaggttag<br>aagccagaggatgacacttc<br>tgcaccataatgagtaccac |

| | |
|---|---|
| | tgttgccatttccaaacagt<br>agtattcattttgctatctt<br>tacctgttcatcttttcgta<br>gtcctaactagcaggctgaa<br><br>ggtaagattactgaggactt<br>ttccaaaagcaaatgtgccc<br>atgtacttatcccatgcata<br>tggcataatcccatttttgt<br>gacgtgaaaaagtctgagct<br>gtattcagcattcccaaaga<br>acagtcatttccttctcgaa<br>gcagtttagcaatatgaacc<br>tgaaaaaatccctccctgct<br>cctcaccagtgagaacacta<br>aagaggggaaggcatcaggt<br>tgtgataaagccggttctga<br>tcggaggagcaaaggttttc<br>gagggaaagcagactcgatc<br>gacaaagcaatgctcacatc<br>tagagatgttcagtcaccag |
| 7-9K | ccacggggaagaaggtttaa<br>ctcctcacacaaacacagat<br>caggaggtgggttctgagag<br>tgtgacaaccaccctcacag<br>gtcttctgaattgaaccctg<br>cattcaggaaacatcagcct<br>tcagggacaagcaacaacca<br>ctcagatggggaaatggaga<br>ctgccaaacatctacattca<br>agagctcaacaaggagtagc<br>agaggcaggagagttcactg<br>aggaagtggctagacctgac<br>cacagaaaagggagaggggc<br>tacatgaaaggtccagcgcc<br>gaatatacatgctgattccc<br>agtgtagaaacagaccagac<br>gccggtacagggaaatacta<br>agcctcttcaaatgtgtttg<br>ggctacatatcctttagtat<br>cagtttaacagctcaagaga<br>ctcacagctgcacgatttaa<br>aggggatacatgtgtacact<br>aaaccataagtgctcactgc<br>ttgagtctgtgaagagagtt<br>aagtgtagctttctggagtc<br><br>gttagtggctatgtagggaa<br>gaaagtagaatggggttgct<br>actgatatgaagtgcttgga<br>gccagtcacagacagacaaa |

| | |
|---|---|
| | aggacactgtgctcagagaa<br>gctacaacatagacgcatct<br>gtccactaatggatgaatgg<br>gatgtgtgtacattacagca<br>gaactgaaggtaggatctca<br>tagtaataccacttctggct<br>actacagtggttcctcaata<br>aatggtgcagttgctatgaa<br>ttgtgcactcttggtgagaa<br>actcctacaagtcaacaaca |
| 9-11K | tttcacatggttagtggtca<br>ccttcccaatgatactaaga<br>ggtttacaatcataagtggc<br>gtgctttttgcaccaacaat<br>ctgatttttaaagtcctggt<br>ggagtacatacatgtttagc<br>tggagaggagtgttcattgt<br>accaagacagtatttagtca<br>tattgactgacctggaacga<br>atacacacacacacacacat<br>accccaaaagccattgaaat<br>tctatatgaccaaaagccat<br>tcagacttcactactgtaga<br>aggtgaccggtgcagtaaaa<br><br>ccacatattgggtacaatgt<br>gaggggggcaaggtataaaa<br>agactcaaagtggggagggt<br>agagtgacataatggacact<br>gtgagagctaacttatgggt<br>ccaaatactgtgtgttctca<br>gtgaagcaactcaggaatgg<br>tggatggagctggaggatat<br>gttttttttttttttgcagc<br>aatgtggtatctctacacca<br>catcaaccagtgagtggata<br>attgcaaagatgtggaacca<br>agacacttgatggcagtaca<br>tgggtatctacccaaaggaa<br>atttgatctagcaatcccac<br>actggagcctaaaatttcca<br>gatcattacattccattgca<br>cctccatttaccagatttta<br>attcttgccttacaggagtg<br>tttgagttccaacagagaga<br>ttgagatttacccagttctg<br>tgtgtcctgaatcaaagatt<br>agcaagtaaaagagagccat<br>attcagaaatgcttaccaca<br>gtcctagtttttgtattgga<br>accataggagttactgtctg |

| | |
|---|---|
| | agtggagaggagtgttcatt<br>gctatttcaatttcatcagt<br>atattgacagacctggaaca |
| 10-12K | ccacatattgggtacaatgt<br>gaggggggcaaggtataaaa<br>agactcaaagtggggagggt<br>agagtgacataatggacact<br>gtgagagctaacttatgggt<br>ccaaatactgtgtgttctca<br>gtgaagcaactcaggaatgg<br>tggatggagctggaggatat<br>gttttttttttttttgcagc<br>aatgtggtatctctacacca<br>catcaaccagtgagtggata<br>attgcaaagatgtggaacca<br>agacacttgatggcagtaca<br>tgggtatctacccaaaggaa<br>atttgatctagcaatcccac<br>actggagcctaaaatttcca<br>gatcattacattccattgca<br>cctccatttaccagatttta<br>attcttgccttacaggagtg<br>tttgagttccaacagagaga<br>ttgagatttacccagttctg<br>tgtgtcctgaatcaaagatt<br>agcaagtaaaagagagccat<br>attcagaaatgcttaccaca<br>gtcctagtttttgtattgga<br>accataggagttactgtctg<br>agtggagaggagtgttcatt<br>gctatttcaatttcatcagt<br>atattgacagacctggaaca<br><br>atctcagacttcactactgt<br>ggaccacataagtggaaggt<br>atgattattaccttggccta<br>gcaagtgagagataaaccca<br>ttagatcagattcttttgca<br>gttccataagattacatgca<br>gctgaatcttaccatggtat<br>taaatgaactatcccctggg<br>tgagctctatccagtaaaca<br>ctaactgaggtatagtcaca<br>ccatggatgggagaactgat<br>aggacactgagaaacagtgt<br>aacacccacatatcacttct<br>ggaagtgaataaggcgaggt<br>ccatgcaacacgtttttcag |
| 12-14K | tatatggagcgtcgtagagg<br>gatggcaactagcagaaact<br>actgcatacatgatagatca |

| | |
|---|---|
| | ccaattacagtcttagtagg<br>gccatggacttacctaaatc<br>catccattacaatagcacat<br>aaattcatacactccacagc<br>catggcctttttctatagtg<br>gtccgggattatagaggaac<br>gttgtcattagtagttatct<br>catccacagccagaaaatgt<br>atgacatgttatagagccat<br>aacctcaaagccatcactag<br>cacactttgagatggctagg<br>agggatgcaaaagagatccc<br>caggtgaagacacctacaga<br>agtgtcaccaaccagatgtg<br>ctttcaacccatctcacaaa<br>tattctattttggcagcacg<br>cacatgaggagaggaccaac<br>cagccactacgcagttttac<br>tgttatactcgattccggta<br><br>tactgttttgtgctaagcca<br>gtaggggaaagaataccagg<br>cacagacgctttcctagaat<br>ggatgccacttaagatttca<br>ggccctttagaagacagaaa<br>tatgaggtaagagccagagc<br>ctgtgtacaatgttctgctc<br>taagtccaccgaggcagatg<br>actcaggtatttccttctag<br>taacagggacagcacaccag<br>cagttgtcaccttgacagaa<br>gacacacagatgatgggtca<br>ctagcaaagcatttgccttt<br>caggtatggctttgaacagt<br>ccacagggtttgaagttcaa<br>gagaaagatgccactgaatc<br>aagacaggctgtgtggtgta<br>gtaactcaacagtccacagt<br>tacagatatgcggtgcttac<br>attcaaacacggctaccaca<br>tggttccccaaaaagggaat |
| 14-16K | catgtacacagatatgcggt<br>aatttacacatgctgtgccc<br>gggggtgtccatcatattta<br>ggtgaaggtgtggggaaatg<br>actattttccacttctcaca<br>atcagtaagtatacaaccca<br>ctcacaaaatccaggaacca<br>gctggattagccttttcaaa<br>gttaggagaaaacatggggt<br>aagtctagatagagtcctcc |

| | |
|---|---|
| | cctgtcacttgttatgctaa<br>ggtgtaaggacaacaggcta<br>gttaatagctgtattagcca<br>aataatgaggtagggcttcc<br>cggcatgaagacatcacagg<br>gacgtaccttattcttgcat<br>aatagacgtgagtggatgga<br>gctttctgtgtaatgcaaag<br>cgaggtagacagaccaagac<br>ggcagtgaggacaactagat<br>atggggtagtcagtcagatg<br><br>ccattaggcagagcaaaacg<br>tcggcgtgacagtatcaagt<br>gcaaagaccactgaactgga<br>tgtgtacgggagcatttcaa<br>acttactggcatttttaaca<br>ggtgcaagctgggtttgtat<br>ctacgctctctgaatgtgac<br>tggaaaacaacccatcccag<br>gaagagctagccacacagtg<br>acattccgggtacacagaac<br>gctggacactagaacaggac<br>agcaggaataggctgttgag<br>aaaaagttccagcaggctgt<br>ccagcatggcaacatatttt<br>catgaaaaaggctgccaggg<br>taggacaattaccccttttgg<br>aactcatcttacagaccacc<br>atgtctattagcaagtcacc<br>gccactgaatagaacatttc<br>tagtgtcgatggttgcacaa<br>cgtctgtttgggatgacgaa<br>gagacaggtaggaagttatt<br>aggttaccaaagactagggg<br>gcacaacaaagtttaccagt<br>ggcaaattcacacagacaga<br>attacacttatatgaggccc<br>agtgaaatcagacagaccca<br>gaatgaaccctgaaaacccg<br>acgcagtactgttatatgct<br>gtggaatattattcagccag<br>acaaaatggggtctatccat |
| 16-18K | accattgttcataatagcca<br>acttgtacgcaactgttcat<br>attctactcctaggtatata<br>atacagttaccatatgaccc<br>tagtttggcagttcctcaaa<br>gaaatggtgcatggacgtgg<br>ctcttacttactggtgggaa<br>gttggcaagaatgcgcagaa |

| | |
|---|---|
| | ggctacttcacactcattag<br>tgcaaatccaaatgacacca<br>atagcatcactcatcagaga<br>aacagcaccagcaaaggtgc<br>cagggaacatacacatatgg<br>aggctcagcacagacacttc<br>cgcctaattaaaaagtgggc<br>agggtctagaatccagaata<br>tttgcaaaccatgtatctga<br>caacccacagaatgggagaa<br>cttgcacagccaaatcaagt<br>aagtgctgggattacaagcg<br>acaagttttcgctatgctgg<br>accatgcctgactaattttt<br>cctgagaagctggtattaca<br>ctgggttcaagtgatcctcg<br>atgatctcagctcactgcaa<br>caaagtctcattctgttgcc<br>tcccttaaatcaagtcactt<br>ctgactgaatatagccagtc<br>aacaccttgagactttttgc<br>cccaagaccttaatcattca<br><br>tttctgtgcctctaaacacc<br>attaagaacccaactgcacc<br>agttgtgcttctcatcattt<br>tgtctgctgacattctacac<br>gatccagcacatctagcagg<br>acaaatggaaatgctttccc<br>caccacacccatcataaatt<br>cacagtgctgggactacagg<br>ggctgatcttgaactcttgg<br>tagggtatcactatgttgcc<br>tttgcataaatccagtacct<br>ctcatctcaaaagacgcaga<br>gggacttagtattccattaa<br>agcttggttcagaaatcaca<br>aataaaactcaggcctagga<br>gcttggccttctaatttatt<br>actaggactacaggcacatg<br>aagcaatcctcttacctcag<br>agtggcgcaatcaaagcttg<br>tttgagacagcgtcttagtc<br>gttttttttttgtcgttttt<br>ccttagtttattttttctgt<br>catttgccagggactgtcat<br>atttgctttacaagtcctgc<br>ctcccaatcttatttgcttt<br>gaactcctgacctcgtgatc<br>aagacggggtttcactatgt<br>cacctggctaattttttttgt |

| | |
|---|---|
| | caagtagctgggactacagg<br>ctgggttcgagattctcctg<br>tctcagctgactgtaacctc |
| 18-20K | caggctggagtgcaatggtg<br>ttgagacggaatcttgctct<br>tcttcttggttttttttttt<br>agaagttctaattcattcct<br>atgatgcccttaagtccaag<br>tgctatcctacccaaaacct<br>cacactcatgaccactgacc<br>accaagtttttgggctcaaa<br>actgagtggcaaagtaggga<br>aatttggccaaagtcatgca<br>ccggaaacttgcttcatgaa<br>tttttatccccatttcattc<br>ccaacggataggtttgaaca<br>attcatttcagtttcacaca<br>tcacaagtacctgcacgata<br>agggacaggaaacagcagct<br>gtctagatatttcccatcat<br>cagtgcagagggttgggaac<br>cagctggtgttacaagaccc<br>agctcctaaaagaaggccag<br>ccccacatctactaaaacaa<br>cagtccgggcaacacagaaa<br>ggatcacttgagtccagaag<br>aacactttgggaggttgagc<br>agtggctcacacctgtaatc<br>aaaagttttcaaggctgggc<br>atttaatagcccagtaaccc<br>tgctactctacactaataca<br>ctgctttttttcaaaaggcca<br>gcagaactgcttactttatc<br>acaaaatccaaggcatgccg<br><br>acaatgcgagcctctagaaa<br>ggcaagcgtgatcataagtg<br>ccctcttccattcttaaaag<br>tagagcaagactgcctcaca<br>caattgaggaaccagcgtgg<br>tcctcggatggcatcagtag<br>gcccacacgaaaccttacat<br>ctctgtggaatgaggcaaca<br>gatgttgatggagtgaccat<br>ctgtcagctgaatgcctgtg<br>agaacagcagcagcgtgaag<br>aatgatttgtgcattgccac<br>ccttggtaactgtcacaact<br>gttagggctgttttcactgg<br>cttcgcagacaggaatgctg<br>cagcttgcatgacttcaggc |

| | |
|---|---|
| | tcacgtcctgaatcctcatg<br>tacagatggaaggccttcat |
| 20-22K | cacttcctcagataaccatt<br>tcaatgaagaaagggagcca<br>atgtgacagagaatcccttc<br>aatggggaggaactgatcca<br>gcagatcggatttgaccaac<br>ctcacttgtcattacacagg<br>aggagtgacggtgagaatgc<br>ctcaggggaagcggagagac<br>catggagcagactgctgagc<br>tgggtcaccctgcatcaaaa<br>ggcgttagtgtcgggaatac<br>agacacagtgaagcccaagc<br>ttaggtctgcactgtcaaca<br>gtccagtgcaaacaagaaga<br>gcagttgaaaataaccctca<br>ccacatagtaactaccaatc<br>ttcagcccaagtaattgtgt<br>ccaaccacaacaggtggatt<br>aaccacaaaatgctggtgcc<br>tagaaggggatttgcccaaa<br>ggttgcaaaccaaaagctgc<br><br>gagctacgcggggaaacgtg<br>gggtgtgtgctgtgggataa<br>ttcagcgtgttagcaacaga<br>gggcagtcaaaacacactct<br>ccttacaaggcctcagaaat<br>gccgatgaagcaacaaagct<br>cattcacggccatctaagtc<br>aaacacactatggtgcgggc<br>gaatgtgttgggccaactgc<br>tcagacctctcaaaggggtg<br>gaatctgagcagaatcagcc<br>agtgctaaggagctcagaag<br>caggacccaaaggtacacaa<br>tcctgtcacttgactctaac |
